# Efficient trace reconstruction in DNA storage systems using Bidirectional Beam Search

**DOI:** 10.1101/2025.04.16.644694

**Authors:** Zhenhao Gu, Hongyi Xin, Puru Sharma, Gary Yipeng Goh, Limsoon Wong, Niranjan Nagarajan

## Abstract

**Motivation:** As DNA data storage systems gain popularity, the need for an efficient trace reconstruction algorithm becomes increasingly important. These algorithms aim to reconstruct the original encoded sequence from its noisy sequenced copies (or “traces”), enabling a faster and more reliable decoding process. Previous works have often been adaptations of methods for multiple sequence alignment or read error correction, typically operating under strict assumptions such as fixed error rates. However, such methods demonstrate limited generalizability to real datasets with higher error rates and suffer from slow processing times when dealing with a large number of traces.

**Results:** We propose a new probabilistic formulation of the trace reconstruction problem. Instead of optimizing alignment among traces, we model the traces as observations of a *k*-th order Markov chain and try to predict the sequence that is generated by the Markov chain with the highest probability. Such a formulation inspires a novel solution, i.e. Bidirectional Beam Search (BBS), whose reconstruction phase operates in linear time with respect to the length of the encoded sequences. Experiments on multiple public Nanopore datasets demonstrate that BBS achieves top-tier accuracy compared with the state-of-the-art methods while being ∼20x faster, showing its potential to enhance the efficiency of DNA data storage systems.

**Availability and Implementation:** The implementation of BBS is available at https://github.com/GZHoffie/bbs, and the dataset and scripts for reproducibility are available at https://github.com/GZHoffie/bbs-test.

## Introduction

DNA data storage has emerged as a transformative solution for modern information storage thanks to its extraordinarily high data density (up to 455 billion GB per gram) [6], long-term data integrity, and low energy consumption [12, 41]. In this paradigm, digital data is encoded as nucleotide sequences, synthesized into DNA strands, and later retrieved through PCR and sequencing [32]. The trace reconstruction problem, which aims to reconstruct the original data from the error-prone sequenced reads, or “traces”, is vital in building an efficient and reliable DNA data storage system [22]. An ideal reconstruction algorithm must be fast to reduce latency, accurate to prevent data loss or resequencing, and be able to work with as few traces as possible to eliminate the need for multiple PCR rounds and deeper sequencing coverages.

Recent advances in third-generation sequencing technologies, notably Oxford Nanopore’s portable sequencers, have expanded the capabilities of DNA data storage [7, 39]. By supporting real-time sequencing of longer read lengths, higher sequencing speeds (*<* 0.02 seconds per base per pore [25], compared to *>* 2 minutes per base in Illumina sequencing [18, 19]), and PCR-free protocols [34], Nanopore technology offers the potential to significantly increase data density and reduce read latency. However, these benefits are offset by a higher mean error rate of around 6% [11], which, along with errors from cost-effective synthesis methods [1], greatly complicate the reconstruction process. Additionally, unlike genome assembly and multiple sequence alignment (MSA) problems, DNA storage systems can leverage prior knowledge such as the length of the encoded sequence and error correction codes (ECC) to aid in reconstruction [30]. Effectively integrating this prior knowledge while mitigating errors in the reads presents significant challenges to the trace reconstruction algorithms.

A large swath of algorithms have been proposed to solve the trace reconstruction algorithm. Based on the various optimization goals, the methods can generally be categorized into alignment-based, IDS-channel-based, assembly-based, and deep-learning-based.

Several popular algorithms used for MSA have also been used for the trace reconstruction problem. For example, Antkowiak et al. utilized the alignment results of MUSCLE along with a weighted majority voting to find the consensus sequence [1, 13], and Xie et al. demonstrated superior error-correcting capabilities using MAFFT [20, 40]. MSA algorithms have also been shown to have good performance under high error rates [1]. However, finding global optimal scoring metrics for the alignment under different synthesis and sequencing conditions is not trivial and has to be determined empirically. In fact, it has been demonstrated that the error situation is highly contextual in Nanopore sequencing and a global error scoring mechanism is often inadequate at encapsulating the nuanced error patterns in Nanopore reads [28]. Moreover, due to the exponential time complexity as a function of the number of sequences, MSA algorithms are limited to small clusters of reads for efficient decoding and cannot handle elevated sequencing depths.

Insertion-Deletion-Substitution (IDS)-channel-based methods model the DNA synthesis and sequencing process as a communication channel, where insertions, deletions, and substitutions occur at each sequence position with probabilities *p*_*I*_, *p*_*D*_, and *p*_*S*_ respectively [36]. These methods aim to reconstruct the original sequence by finding the consensus string that maximizes the likelihood of observing the given traces [4] under the aforementioned assumption. In practice, this optimization is typically performed by iteratively correcting errors in the reads until they converge to a consensus, as seen in the Bitwise Majority Alignment (BMA) algorithm [3]. Several variants of BMA have been proposed, including BMA Lookahead, which improves error correction by looking at a longer range of bases in the reads [15, 23, 27], and Trellis BMA, which refines the IDS-channel with a hidden Markov model [36]. Although these approaches are theoretically well-founded under the IDS-channel model, they may face challenges when applied to real datasets, where the assumption of independent error occurrence does not always hold. This is especially evident for third-generation sequencing technologies, such as Nanopore, where base-pairs are not individually sequenced but rather DNA fragments are indirectly deduced by decoding the electrochemical signals of consecutive overlapping 10-to-20 base-pair oligo fragments. Consequently, successive errors are not independent and sequencing errors are often more prevalent at the two ends of a read than the middle section [1, 24], leading to deviations from the expected error model. Similar to multiple sequence alignment (MSA) algorithms, building a universally robust error model remains a challenge.

While previous methods depend largely on the chosen hyper-parameters and/or the error model determined using the training data, the assembly-based methods offer a “parameter-free” approach, aiming to maximize the number of reads that agree with the subsequences of the final consensus. This is achieved by greedily identifying the maximum-weighted path in the de Bruijn graph using DBGPS [35] and by assembling the longest common subsequences between each pair of reads in the iterative algorithm (ITR) [31]. These methods demonstrated superior accuracy across different datasets. However, they are computationally intensive and fail to fully account for the prior knowledge of the encoded sequence.

Recently, deep-learning-based methods emerged as a popular alternative for trace reconstruction, which optimizes the alignment and consensus construction process by auto-learning the error patterns. RobuSeqNet assigns weights to each read using convolutional layers and attention mechanism [29], and finds the consensus sequence based on the weighted sum of all sequences within the cluster. DNA-GAN [42] and Single-Read Reconstruction (SRR) Algorithm [26] attempted to output the consensus sequence directly in the network output. Such methods show promising results but incur high computational costs and require special hardware. Furthermore, they might not perform equally well given unseen error distributions.

In this work, we aim to propose a computationally lightweight yet highly accurate trace reconstruction algorithm that is capable of reproducing the original data sequence at low sequencing depths. Our work extends both assembly-based and deep-learning-based methods for trace reconstruction algorithms. Rather than optimizing sequence alignment or learning a global error model for the entire dataset, we model the traces within each cluster as observations from a *k*-th order Markov chain and aim to predict the sequence with the highest likelihood of being emitted. This approach allows error probabilities to be estimated independently at each position within every strand cluster, providing greater flexibility in handling complex sequencing errors.

To efficiently identify the most probable sequence, we introduce the bidirectional beam search (BBS) algorithm, which leverages the learned Markov chain to determine the most likely next trace. Notably, the computational complexity of the reconstruction phase of our BBS algorithm scales linearly with the length of the consensus sequence, making it highly efficient. Experimental results on multiple Nanopore datasets and various cluster sizes demonstrate that our approach is among the most accurate methods when compared with the state-of-the-art algorithms while being ∼20x faster, highlighting its potential to improve the efficiency of DNA data storage pipelines significantly.

## Method

### Problem Formulation

Let us denote the set of alphabets to be Σ = `{A`, `C`, `G`, `T}`. Algorithms for trace reconstruction problems take a set of *N* traces, *C* = {*c*_1_, …, *c*_*N*_}, as input, where each *c*_*i*_ in *C* is a string Σ^∗^ that is some erroneous version of a seed string *s* ∈ Σ^*L*^, and aim to output the seed string *ŝ* such that *s* = ŝ with high probability. For simplicity, we denote *S*_*i*:*j*_ as the subsequence of *S* from the *i*-th to the *j*-th character, *S*_*i*_, *S*_*i*+1_, …, *S*_*j*_. We also write 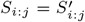 to represent the event 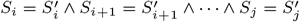.

While the intuition and the purpose of the trace reconstruction problem are clear, the formal definition of the optimization objective varies across different methods. MSA-based methods attempt to find the consensus in the alignment of all the traces that maximize the global alignment score [1, 13, 20, 40], while deep-learning-based methods minimize the loss function that captures the reconstruction errors measured using edit distance [35, 42]. The most widely used definition is to model the synthesis and sequencing process as an insertion-deletion-substitution (IDS) channel and attempt to find the seed string that maximizes the likelihood of observing the traces [4, 36].

**IDS-channel-based trace reconstruction.**

**Input**: A set of traces *C*.

**Assumptions**:

1. Each trace *c*_*i*_ ∈ *C* is independent.
2. Each trace *c*_*i*_ is a modified version of the seed string *s* ∈ Σ^*L*^, where insertion, deletion, substitution, and matching happen at each position with probability *p*_*I*_, *p*_*D*_, *p*_*S*_, *p*_*M*_ respectively, with *p*_*I*_ +*p*_*D*_+*p*_*S*_+*p*_*M*_ = 1.

**Output**: arg 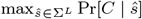.

Under the assumptions, the log-likelihood of observing the traces can be calculated by

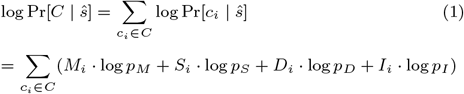

where *M*_*i*_, *S*_*i*_, *D*_*i*_, and *I*_*i*_ denote the total number of matches, substitutions, deletions, and insertions needed to transform ŝ to the trace *c*_*i*_. The goal can then be interpreted as finding the length-*L* string ŝ with the highest alignment score defined by *p*_*I*_, *p*_*D*_, *p*_*S*_, and *p*_*M*_ to all the traces.

This problem definition, though supported by many theoretical works [8, 9, 10, 17, 21, 37], might encounter problems when applied to real-life data. Firstly, solving the optimization is hard. In the general case, the best existing solution to the optimization problem is brute-force [4], taking exponential time with respect to sequence length *L*. Secondly, even if the optimal string *ŝ* is found, it doesn’t necessarily reflect the true alignment. In real datasets, the fixed *p*_*I*_, *p*_*D*_, *p*_*S*_, and *p*_*M*_ can vary across different clusters and positions within the traces. Notably, error rates tend to be higher at the ends of reads [1], leading to inconsistencies and reducing the reliability of alignment scores. Choosing the correct values for *p*_*I*_, *p*_*D*_, *p*_*S*_, and *p*_*M*_ is challenging since different definitions of these parameters can result in drastically different alignments and consensus sequences. Furthermore, when incorporating prior knowledge of the encoded sequence length, the optimal alignment may not accurately represent the true consensus, as illustrated in the example in Figure 1.

**Fig. 1:**
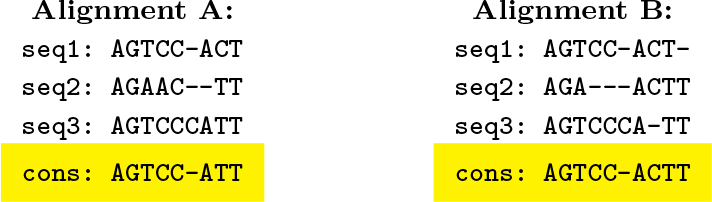
The optimal alignment and consensus (denoted `cons` in the figure) may differ given different scoring metrics. Neither of the alignments is correct if given the prior knowledge that the length of the encoded sequence is 10, in which case the optimal consensus would be AGTCCCACTT.

Therefore, we propose an alternative objective. Instead of finding the seed string that maximizes the likelihood of observing the traces, we model the traces as observations of a *k*-th order Markov chain (*k*-MC) consisting of *L* variables, *S*_1_, …, *S*_*L*_, where *S*_*i*_ ∈ Σ = `{A`, `C`, `G`, `T}`, and assume that for all *i* = *k* +1, *k* +2, …, *L*,

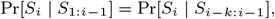

and try to predict the most probable trace that is generated by the *k*-MC. We can then learn the conditional probabilities *𝒟*_*C*_ from the set of traces *C*, and perform the maximum likelihood estimate (MLE) of the future trace.

***k***-MC-based trace reconstruction.

**Input**: A set of traces *C*.

**Assumptions**: Each trace *c*_*i*_ ∈ *C* can be viewed as independent observations of a *k*-th order Markov chain.

**Output**: 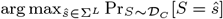.

Under the *k*-MC assumption, the joint probability of observing *𝒟* as the next observation can be calculated by

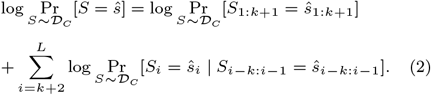

Intuitively, the new problem definition also tries to find the underlying seed string used to generate the traces. Under the IDS-channel assumption, if *p*_*M*_ = max{*p*_*M*_, *p*_*I*_, *p*_*D*_, *p*_*S*_}, the most probable future trace given a seed string *s* is *s* itself, with the highest likelihood of 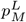.

The advantages of the new problem definition lie in the following: Firstly, exact as well as heuristic algorithms to find the most probable observation in a Markov model, such as the Viterbi algorithm, are well studied [38]. Secondly, it uses weaker assumptions than the previous definition based on IDS-channel, which is essentially a first-order hidden Markov model with fixed probability values for errors. Since *k* is chosen such that all the (*k* + 1)-mers are unique in the sequences, the conditional probabilities 𝒟 _*C*_ for each cluster *C* allow us to learn different error probabilities for each position in each cluster. Thirdly, calculating the likelihood for the next trace doesn’t require any alignment or calculation of edit distance among the traces, leading to high efficiency. Finally, the value 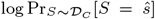 represents the joint likelihood of observing *ŝ* as the next trace, allowing us to assign a confidence score to the output, as well as compare the goodness of the output sequence, aiding future post-processing and the downstream decoding tasks.

Next, we propose a novel solution, i.e. bidirectional beam search (BBS), to efficiently find the most probable observation of the estimated *k*-MC with high probability.

### Bidirectional beam search (BBS) algorithm

The main idea of the bidirectional beam search algorithm is to incorporate the learned parameters *k*-MC model into the de Bruijn graph and find the best path that represents the most probable sequence generated by the *k*-MC model using beam search. The beam search is done in two directions, one on the original traces and one on the reversed traces, for optimal performance. An illustration of the BBS algorithm is shown in Figure 2.

**Fig. 2:**
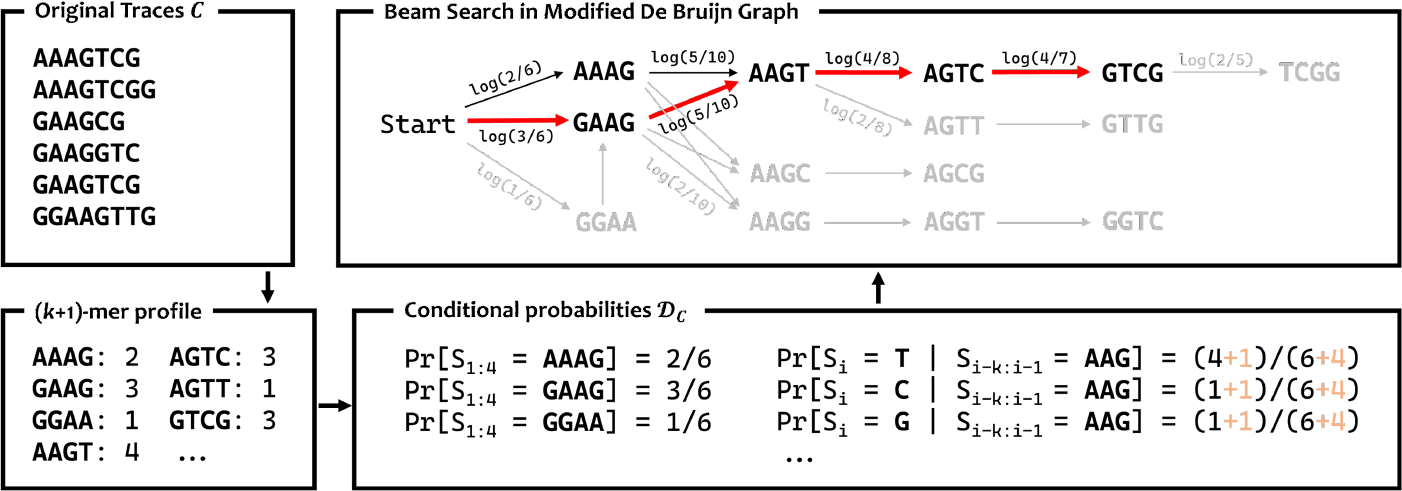
The general workflow of the BBS algorithm. In the first step, the set of original traces *C* (top left) is converted into a hash table that maps each (*k* + 1)-mer to the number of times it appears in *C* (bottom left). *k* is chosen such that all the *k*-mer appear at most once in each trace. We then use the (*k* + 1)-mer profile to estimate the conditional probabilities 𝒟 _*C*_ (bottom right), which replaces the edge weights in the De Bruijn graph (top right). Finally, beam search (with beam width *B* = 2 in the figure) is performed to find the best *w* paths of length *L* in the De Bruijn graph. It saves time by skipping paths that are not promising (marked gray) in the graph. For clarity, only one directional beam search is shown in the figure.

The first step of the BBS algorithm is to choose the value of *k*. In our algorithm, we choose the smallest *k* within a predefined range [*k*_min_, *k*_max_] such that all the *k*-mers are unique within each trace. The expected value of *k* is *O*(log *L*) (**supplementary material section 3**). In rare cases where such *k* cannot be found, *k*_max_ is used. Such cases can be avoided by carefully designing the encoding scheme in practice. The uniqueness ensures that the de Bruijn graph built using the (*k* + 1)-mers in the traces is a directed acyclic graph (DAG). We also choose the smallest *k* that satisfies uniqueness as smaller *k* can lead to better predictions of the conditional probabilities and, consequently, a better reconstruction performance. We then store the (*k* + 1)-mer counts in a hash table. This process takes time linear to the size of the input.

We can now build a modified de Bruijn graph with the following specifications,

- **Vertices** (denoted *V*): All possible (*k* + 1)-mers that appear in the traces, plus a special vertex `Start` indicating the start of the reads.
- **Edges** (denoted *E*): For each (*k* + 1)-mers *v* ∈ Σ^*k*+1^ that have appeared at the start of each trace, a directed edge (`Start`, *v*) is added. We also add a directed edge between pairs of (*k* + 1)-mers (*u, v*) if the last *k* bases of *u* matches the first *k* bases of *v*. Note that this does not require adjacency of *u* and *v* in the same sequence.
- **Edge weights**: For each edge (`Start`, *v*) ∈ *E*, we assign the weight to be the estimated joint probability of the first (*k* + 1)-mer in *s* being *v*,

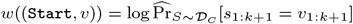

while for other edges (*u, v*) ∈ *E*, we assign the weight to be the estimated conditional probability.

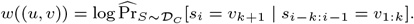

Based on the *k*-MC assumption, we can estimate the joint probability and the conditional probabilities using the maximum likelihood estimate,

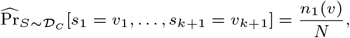

where *n*_1_(*v*) refers to the number of times the (*k*+1)-mer *v* appears as the first (*k* + 1)-mer in the traces, while

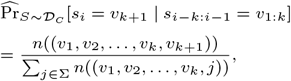

where *n*(*v*) = *n*((*v*_1_, …, *v*_*k*+1_)) refers to the number of times (*k* + 1)-mer *v* appears in anywhere in the traces. In practice, especially with a high error rate and small coverage, the numbers *n*(*v*) can be small, resulting in inaccurate estimations. We therefore apply Laplace smoothing,

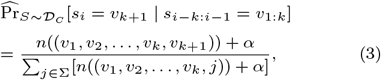

and set the default *α* = 1. Intuitively, larger *α* would lead to larger penalties on “rare” paths that very few traces agree.

The construction is similar to the typical de Bruijn graphs used for assembly except for the assignment of edge weights. With this construction, a path starting from `Start` vertex and containing *L* − *k* + 1 vertices would correspond to a string *s* ∈ Σ^*L*^, and weight of the edges in the path sum up to 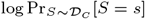. Thus, the trace reconstruction problem is reduced to a fixed-length longest path problem,

#### Algorithm 1

Beam Search for Trace Reconstruction

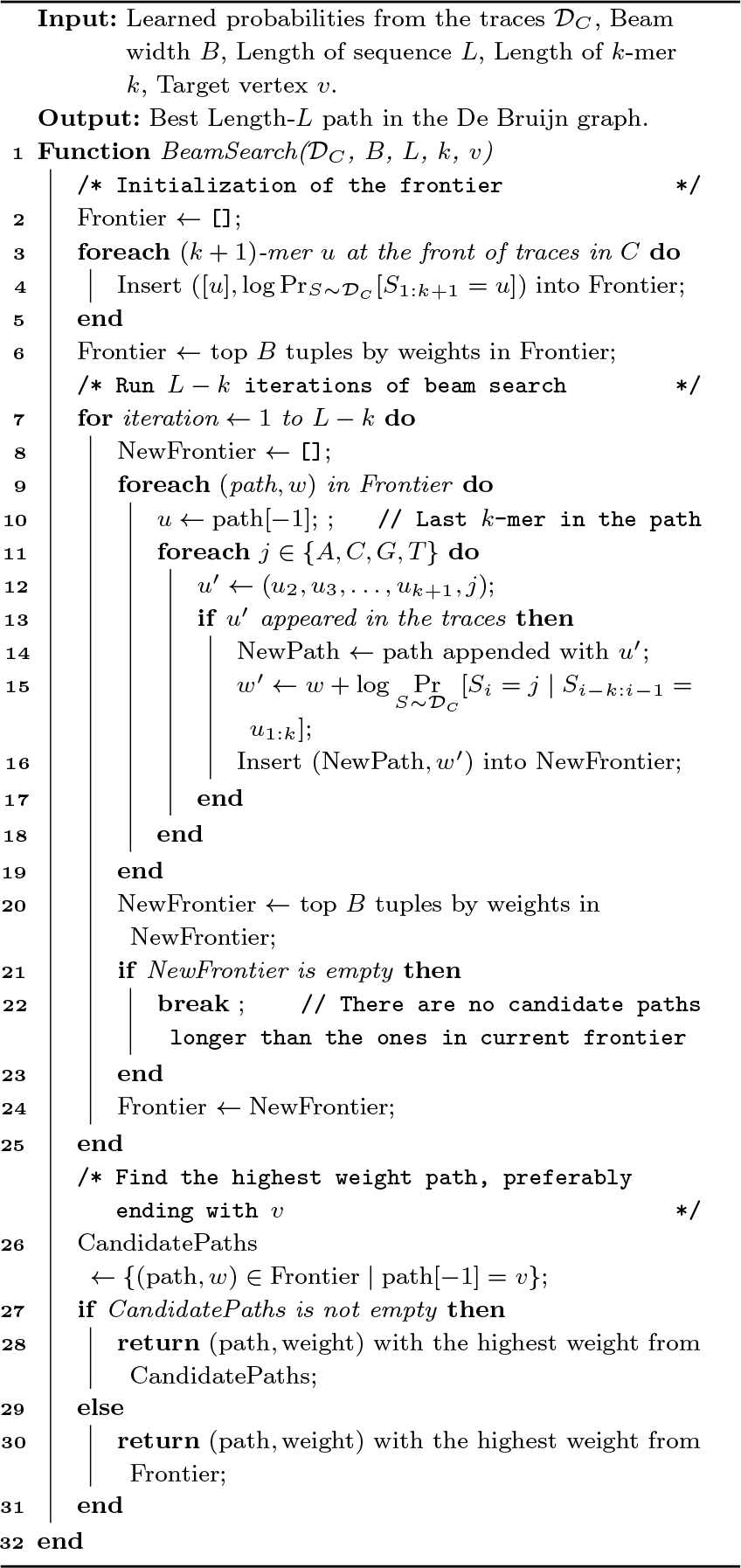

**Fixed-length longest path problem**

**Input**: A directed acyclic graph *G*(*V, E*), the starting vertex *u* ∈ *V* and an ending vertex *v* ∈ *V*, and a predefined length *l*.

**Output**: A length *l* path that starts from *u*, ends in *v*, with the maximum sum of edge weights.

In our constructed graph, the length *l* = *L* − *k* + 1, and the goal vertex *v* are set heuristically to be the most frequently appearing (*k* + 1)-mer at the end of the traces.

A brute-force algorithm to solve the fixed-length longest path problem is to perform *l* iterations of breadth-first search (BFS) that allows repeated visiting of the same vertex. However, such an algorithm may result in *O*(|Σ|*l*) possible paths of length *l* in the end. Therefore, we opt to use beam search to approximate the optimal length-*l* path in the constructed graph. The benefits of using beam search involve its fast running time and its ability to output multiple candidate good consensus sequences, allowing more room for errors. By running exactly *l* iterations of beam search, we also leverage the prior information of the sequence length. A detailed description of the beam search can be found in algorithm 1.

In each iteration of beam search, at most *B* paths are kept in the frontier, and at most *B* ·|Σ| paths are explored. The calculation of conditional probabilities is done during the beam search using equation 3, which costs *O*(1) time for each calculation as the (*k* + 1)-mer profile is stored using hash tables. At the end of each iteration, the top *B* paths with the highest weight are inserted into the new frontier. The top *B* tuples are selected using Hoare’s selection algorithm [16], costing *O*(*B* · |Σ|) time. This bounds the time complexity of the beam search to be *O*(*LB*|Σ|), which can be regarded as *O*(*L*) for small *B* and |Σ|.

Notice that the joint probability in equation 2 can also be calculated in a reversed manner,

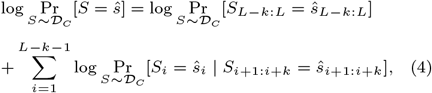

which hints that the beam search can also be done in the other direction that starts from the end of the traces. An intuitive way is to perform a beam search in both directions and select the best sequence with the highest path weight, as shown in algorithm 2.

In the actual implementation, the conditional probabilities 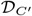 are calculated from the same (*k*+1)-mer profile instead of reversing every trace in *C* to save time. As algorithm 2 essentially performs algorithm 1 twice, the time complexity is also *O*(*L*).

## Results

### Experiment setup

To test the efficiency and correctness of our algorithm, we compare the performance against the state-of-the-art algorithms in each type of method for trace reconstruction. In particular, MUSCLE [13] from the MSA-based methods, TrellisBMA [36] from the IDS-channel-based methods, and ITR [31] from the assembly-like methods. We utilized the latest MUSCLE v5 [14] and implementation by Antkowiak et al. to return the consensus sequence from the alignment. The newest deep-learning-based methods were not included due to the lack of publicly available code [42]. Instead, we compare our results with their reported accuracy.

Additionally, we compare our method against a newly published method, Conditional Probability Logic (CPL), which is a refinement step for the deep learning method DNAFormer [2]. Though BBS was developed independently of CPL, they share a similar methodology. CPL utilizes a first-order Markov chain with the conditional probabilities estimated using the pairwise alignments, instead of *k*-mer counting in BBS, resulting in a time complexity of *O*(*NL*^2^). In addition, CPL finds the optimal longest path in the constructed graph, whereas BBS employs a greedy algorithm for efficiency.

#### Algorithm 2

Bidirectional Beam Search

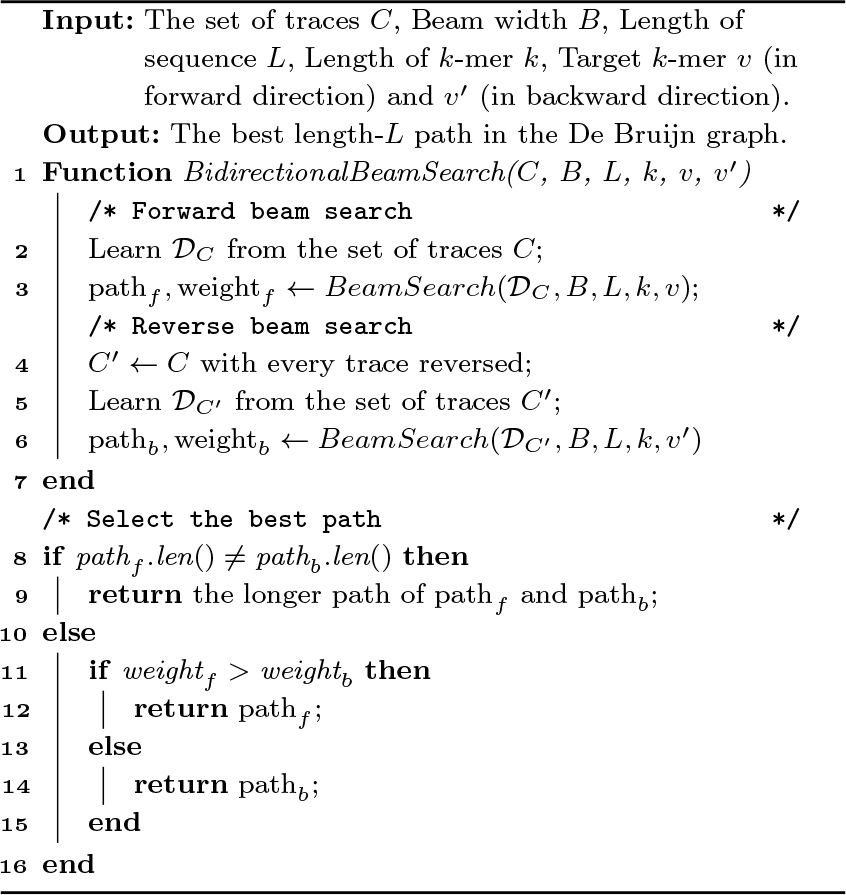

All experiments are run on a machine with Intel Core i9-13900H CPU, with 20 threads and 16 GB of memory. MUSCLE is the only tool with multi-threaded implementation. All tools are run using their default parameters. For BBS, the beam width is chosen to be 20. No error correction code is assumed.

Similar to previous works, we use three metrics to measure the correctness of the algorithms, including the following,

1. Success rate, defined by the percentage of the clusters that are reconstructed correctly without any error.
2. Average Hamming distance per cluster, where the Hamming distance between the ground truth sequence *s* and the predicted sequence *ŝ* is

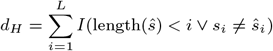

where *I*(*A*) is the indicator function which returns 1 if *A* is evaluated to `true` and 0 otherwise.
3. Average edit distance per cluster, where the edit distance is calculated from the optimal alignment between *s* and *ŝ*.

### BBS allows accurate and efficient trace reconstruction from real Nanopore reads

We test the five algorithms on real datasets, with three open-source datasets published by Bar-Lev et al. [2] (Nanopore two flowcells), Srinivasavaradhan et al. [36] and Chandak et al. [7], as shown in Table 1. The dataset from Bar-Lev et al. is randomly downsampled to 10,000 clusters so that the tested tools finish in a reasonable run time. Clustering for the dataset from Chandak et al. is done by simply mapping the sequenced reads to the closest sequence in the ground truth (perfect clustering). On the other hand, the dataset Bar-Lev et al. uses its own binning algorithm [2], and the Srinivasavaradhan et al. utilize the clustering algorithm from Rashtchian et al. [30]. The performance of each algorithm is listed in Table 2.

**Table 1.**
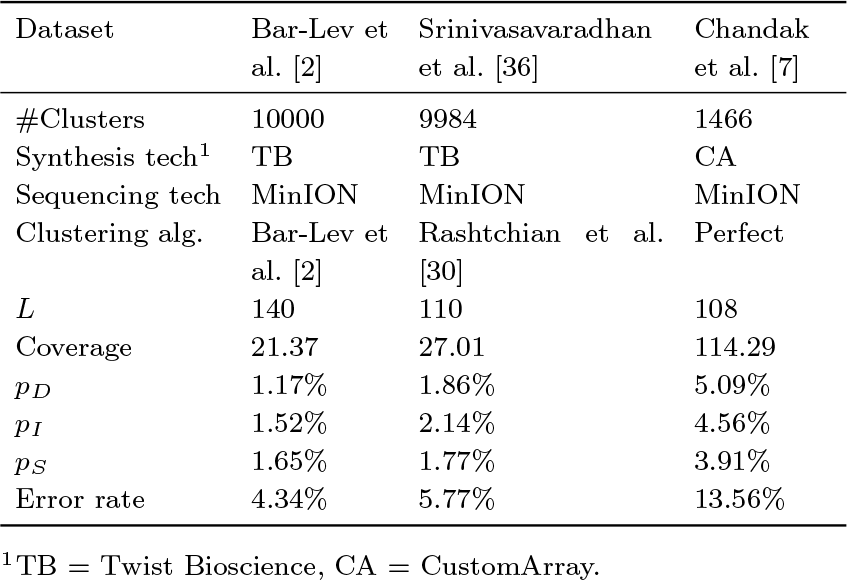
Statistics of the three real datasets. The error rates are estimated using the script by Srinivasavaradhan et al. [36]

| Dataset | Bar-Lev et al. [2] | Srinivasavaradhan et al. [36] | Chandak et al. [7] |
| --- | --- | --- | --- |
| #Clusters | 10000 | 9984 | 1466 |
| Synthesis tech<sup>1</sup> | TB | TB | CA |
| Sequencing tech | MinION | MinION | MinION |
| Clustering alg. | Bar-Lev et al. [2] | Rashtchian et al. [30] | Perfect |
| $L$ | 140 | 110 | 108 |
| Coverage | 21.37 | 27.01 | 114.29 |
| $p_D$ | 1.17% | 1.86% | 5.09% |
| $p_I$ | 1.52% | 2.14% | 4.56% |
| $p_S$ | 1.65% | 1.77% | 3.91% |
| Error rate | 4.34% | 5.77% | 13.56% |

<sup>1</sup>TB = Twist Bioscience, CA = CustomArray.

**Table 2.** Performance of MUSCLE [13, 1], Trellis BMA [36], ITR [31], CPL [2], and BBS (this work) on the three real datasets. The bold text indicates the best-performing tool, while the underlined text indicates the second-best one..

| Dataset | Tools | Success rate | Avg. edit distance | Avg. Hamming distance | Running time (s) |
| --- | --- | --- | --- | --- | --- |
| Bar-Lev et al. [2] | MUSCLE | 96.25% | 0.258 | 1.382 | 1475.55 |
|  | Trellis BMA | 92.41% | 0.363 | 1.635 | 19759.85 |
|  | ITR | 97.42% | 0.221 | 0.975 | 12462.04 |
|  | CPL | <u>98.39%</u> | **0.201** | <u>0.374</u> | <u>495.22</u> |
|  | BBS | **98.80%** | <u>0.206</u> | **0.346** | **23.84** |
| Srinivasavaradhan et al. [36] | MUSCLE | 84.70% | 0.259 | 6.032 | 1741.65 |
|  | Trellis BMA | 69.57% | 0.908 | 6.003 | 17859.17 |
|  | ITR | 87.58% | 0.232 | 4.797 | 7351.52 |
|  | CPL | **94.93%** | **0.150** | **1.286** | <u>361.05</u> |
|  | BBS | <u>94.77%</u> | <u>0.168</u> | <u>1.616</u> | **20.01** |
| Chandak et al. [7] | MUSCLE | 76.94% | 0.366 | 8.821 | 7082.75 |
|  | Trellis BMA | 52.32% | 1.608 | 13.094 | 10744.21 |
|  | ITR | 64.12% | 0.666 | 12.231 | 1614.64 |
|  | CPL | <u>90.93%</u> | **0.205** | **2.299** | <u>131.60</u> |
|  | BBS | **91.34%** | <u>0.283</u> | <u>2.738</u> | **8.52** |

BBS consistently achieves top-tier reconstruction accuracy across all three datasets while significantly reducing the running time. Against MUSCLE, Trellis BMA, and ITR, BBS achieves 3-6x smaller Hamming distance, indicating its ability to reliably reconstruct the encoded sequence, especially for datasets with high error rates and higher coverage.

Notably, the success rate of the BBS algorithm in the dataset from Srinivasavaradhan et al. (94.772%) also surpasses the reported success rate of deep learning methods, such as 85.42% in DNAFormer [2] and 91.73% in DNA-GAN [42], without the need of training and post-processing.

In the meantime, with the same computational resources, BBS is also ∼20x faster than the state-of-the-art algorithms, using only seconds for datasets that used to take minutes to hours to reconstruct. To test the time complexity of the algorithms, we generated clusters of size ranging from 10 to 60 to test the time complexity of the algorithms (**supplementary material section 1**). Both BBS and Trellis BMA scale linearly with the cluster size. However, Trellis BMA shows a much larger constant factor, making it the slowest of the tested algorithms. MUSCLE, being a multiple sequence alignment algorithm, demonstrated an exponential increase with respect to the number of traces. Finally, both ITR and CPL showed a quadratic increase for cluster sizes 10-30 while being roughly constant for larger cluster sizes. This is because ITR reduces the runtime by using only the first 25 traces for cluster size larger than 25 [31]. Such a strategy works fine for small error rates or datasets with low coverage but demonstrates a decrease in performance for datasets with high coverage, such as the Chandak dataset, due to the loss of information.

In addition, we plot the probability that the ground truth sequence *s* disagrees with the reconstructed sequence 𝒟 on the *i*-th position, Pr[*s*_*i*_ ?= *𝒟*_*i*_], against *i* in Figure 3. Both MUSCLE and ITR show a gradual increase in error rate, which is expected because an early wrong decision or an insertion or deletion error at the front of the reconstructed sequence can lead to disagreement in all subsequent positions. Both methods also show a sudden increase in error rate towards the end of the sequence due to the fact that the prior information of the encoded sequence length *L* is not fully utilized. As a result, the reconstructed sequence of these two methods often varies in length. On the other hand, Trellis BMA exhibits a triangular shape since its reconstruction process starts from the two ends and joins in the middle of the sequence. It also shows a larger slope compared with MUSCLE and ITR.

**Fig. 3:**
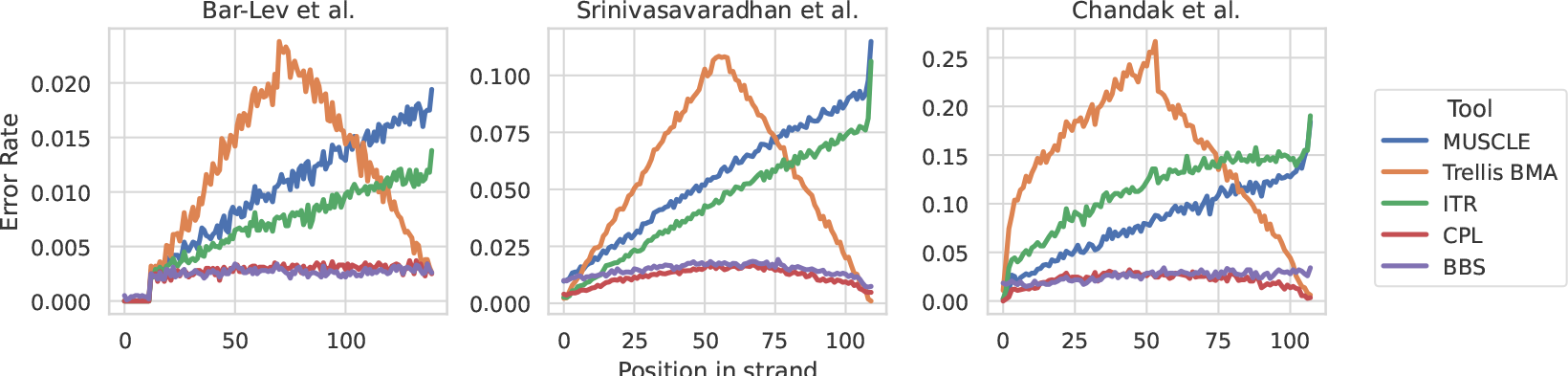
Probability of the ground truth sequence disagreeing with the reconstructed sequence on the *i*-th position, for all 1 ≤ *i* ≤ *L*.

Different from the other methods, the error curves of BBS and CPL are the closest to uniform. BBS chooses the best paths out of all the length-*l* paths, leveraging the prior information on the length of the encoded sequence, and maximizing the global joint likelihood of observing the whole sequence, rather than focusing on local subsequences.

While BBS has among the best error correction performance in real data, its advantage over the other algorithms significantly reduces in synthetic data where errors are assumed to be independent and uniformly random (**supplementary material section 2**). The discrepancy of error correction performance between real and synthetic datasets highlights the limitation of over-simplifying the Nanopore error model, and demonstrates the advantage of the model- and alignment-free approach of BBS.

### BBS achieves top-tier accuracy at all sequencing coverages

Scoring a higher error correction percentage is essential to reducing cost and latency for data retrieval in DNA storage systems. When a trace reconstruction fails, the entire lengthy biochemical process has to be repeated afresh through an expensive re-read. While the accuracy of a trace reconstruction algorithm can be improved by having larger clusters, it involves ramping up the PCR amplification and increasing the sequencing depth, both of which increase the latency and the cost of a data read. Thus, achieving high reconstruction accuracy at shallow sequencing depth is instrumental in reducing the re-read frequency and reducing stuttering in data retrieval while maintaining a low-cost DNA storage system.

To evaluate the performance of our reconstruction algorithm at varying sequencing depths, we randomly sampled 500 clusters from the Srinivasavaradhan et al. dataset [36] and performed a subsampling experiment. For each cluster, we randomly selected a subset of reads to maintain a fixed cluster size. We began by subsampling each cluster to a size of exactly 2, then repeated the process for cluster sizes ranging from 4 to 30. The failure rate, defined as 1 − success rate, was plotted against the cluster size (coverage) in Figure 4.

**Fig. 4:**
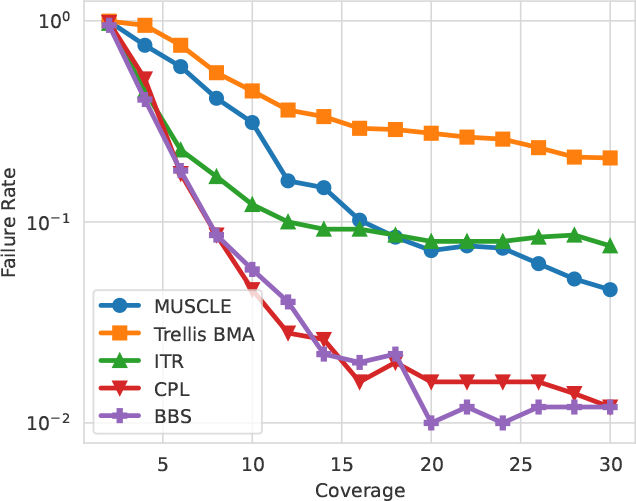
Performance of the algorithms at small coverages in the dataset from Srinivasavaradhan et al.

As expected, all algorithms show improved performance with larger coverage. BBS and CPL consistently outperform the others across all cluster sizes. They also demonstrate the fastest decrease in failure rate, converging to the lowest failure rate. Notably, BBS achieves a success rate of at least 95% at coverage of just 12 reads, while the MSA- and IDS-channel-based algorithms only reach this threshold at a coverage of 30 or higher, striking a 60% sequencing cost reduction for high-success reads. This highlights the ability of our algorithm to perform effectively with smaller coverages, offering the potential to shorten latency and save costs by reducing the required sequencing depth or PCR amplification rounds.

### The path weight returned by BBS is a good indicator of reconstruction quality

Previous non-deep-learning-based trace reconstruction methods only output one reconstructed sequence per cluster with no indicator of reconstruction quality. However, with the *k*-MC model and beam search, we are able to output multiple candidate reconstructions, each associated with a path weight that represents the log-likelihood of observing the sequence as the output of the *k*-MC model. Furthermore, we can also calculate a confidence score for the returned sequence. Let *w*_*i*_ be the weight of the *i*-th path (denoted *p*_*i*_) in the final frontier, the confidence for the *i*-th path can be calculated using a softmax function,

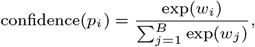

which intuitively represents the probability of the *i*-th path being the output of the *k*-MC, given that the output is one of *p*_1_, …, *p*_*B*_.

We examine the distribution of path weights and the confidence score of correctly and incorrectly reconstructed clusters from the dataset from Srinivasavaradhan et al., respectively (**Figure 5**). It is clear that the correct and incorrect reconstructions result in a very different distribution of path weights and confidence scores. In particular, the incorrect reconstructions tend to have lower path weights and lower confidence scores. The areas under the receiver operating characteristic curve (AUROC) of the path weight and the confidence score are 0.85 and 0.87, respectively.

**Fig. 5:**
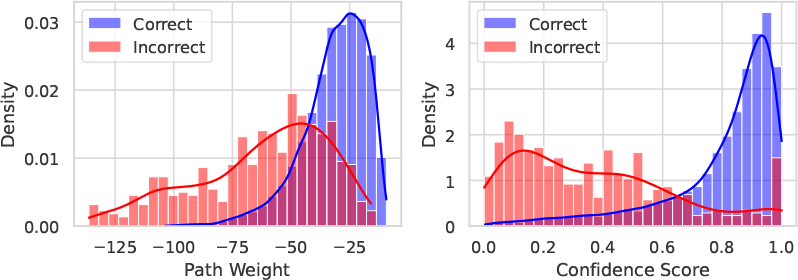
Distribution of the path weight and confidence score for the correctly (blue) and incorrectly (red) reconstructed clusters in the dataset from Srinivasavaradhan et al

We also attempt to fit a simple logistic regression model using the path weight and confidence score to predict whether the reconstruction is correct or not. On the dataset from Srinivasavaradhan et al., the logistic regression model achieves an AUROC of 0.94. This result indicates the potential use of returned path weights by the BBS algorithm to infer the correctness of reconstruction and a hybrid algorithm where more candidate reconstructions from the beam search or more sophisticated but slower algorithms can be used for clusters with low confidence scores, further enhancing the accuracy of the reconstruction.

### Each component of BBS is essential for high accuracy

Finally, we conduct an ablation study to test the performance of BBS with each component removed. Specifically, we apply the same beam search algorithm to a graph where the edge weights are defined by standard de Bruijn graph (DBG) edge weights based on *k*-mer counts. The resulting performance is significantly lower than BBS and only marginally higher than the ITR algorithm (“DBG weight” in Table 3). In addition, the algorithm’s performance also decreases significantly if the Laplace smoothing is not applied (“No smoothing” in Table 3). This shows that our replacement of edge weights with the log of conditional probabilities with Laplace smoothing is well-suited for the task of consensus finding.

**Table 3.** Ablation study results showing the change in success rate on the three datasets when components of BBS are removed.

| Dataset | Bar-Lev et al. [2] | Srinivasavaradhan et al. [36] | Chandak et al. [7] |
| --- | --- | --- | --- |
| **BBS** | **98.80%** | **94.77%** | **91.34%** |
| DBG weight | −0.39% | −4.85% | −8.33% |
| No smoothing | −1.90% | −2.75% | −11.74% |
| Unidirectional | −0.13% | −0.72% | −0.82% |
| Greedy Search | −2.50% | −11.01% | −14.33% |

We also highlight the superiority of bidirectional beam search by checking the performance of the algorithm with a one-directional beam search (“Unidirectional” in Table 3) and a beam width of *B* = 1 (“Greedy Search” in Table 3). The consistent superior performance of BBS over one-directional beam search across all beam widths (**Supplementary material section 3**) further supports the effectiveness of path weight as an indicator of reconstruction correctness. Additionally, the greedy best-first search exhibited a significant drop in success rate, highlighting its reduced flexibility and accuracy compared to the beam search.

## Discussion and Conclusion

The recent surge in the popularity of DNA data storage has prompted the emergence of many algorithmic problems. The trace reconstruction problem, being the core step in the decoding process, is crucial for an accurate, fast, and reliable DNA storage system. While previous methods focused on optimizing the seed string to maximize the likelihood of observing the traces, we propose a new objective to predict the sequence that is most likely the next trace in the cluster. The likelihood can be calculated by modeling the traces as observations of a *k*-th order Markov chain, where the conditional probabilities can be estimated simply by counting the (*k* + 1)-mers in the cluster.

The new problem formulation and model inspire a novel alignment-free algorithm named Bidirectional Beam Search (BBS), which predicts the most likely next trace by replacing weights in the De Bruijn graph with the modeled probabilities in the *k*-MC and finding the longest path with a fixed length with beam search. The beam search is performed once starting from the start of the sequences, and once from the end. The beam search operates in linear time with respect to the length of the encoded sequence.

Experiments on real datasets sequenced by Nanopore show that BBS is uniquely accurate and fast, especially when dealing with datasets with high error rates and large coverage. A simulated study also demonstrated that BBS performs the best for all coverages, achieving the same success rate using fewer traces than the other algorithms. Moreover, unlike the previous methods that only report the reconstructed sequence without any indicator of the quality of reconstruction, BBS reports the path weight along with a confidence score, which proves to be a reliable way to find the wrongly reconstructed sequences, aiding future post-processing and decoding. The experiment results show the potential of the BBS algorithm to greatly enhance the efficiency of the current DNA data storage pipeline.

Possible future directions to improve the algorithms involve optimizing the choice of parameters, including the beam width *B* and the order of the Markov chain *k*. Due to the fact that BBS relies on (*k* + 1)-mer profile for trace reconstruction, the algorithm would inevitably fail if one of the (*k* + 1)-mer in the encoded sequence appears erroneous in all the traces. As a result, a careful choice of *k* is needed for datasets with high error rates and low coverage. In addition, the optimal choice of *B* can also vary with different coverages. Choosing the optimal values can again enhance the accuracy of the BBS algorithm (**Supplementary material section 3**).

It is also worth investigating whether the *k*-MC model can be used in conjunction with error correction codes (ECC) [5, 33]. Though not tested in this work, ECC can be incorporated smoothly into the beam search procedure, where paths that do not satisfy the ECC requirements are filtered out before pushing into the frontier.

## Acknowledgments

This work was supported by the NUS Research Scholarship from the School of Computing, National University of Singapore.

