## Supplementary Material for "Efficient trace reconstruction in DNA storage systems using Bidirectional Beam Search"

### Efficient trace reconstruction in DNA data storage systems with Bidirectional Beam Search - Supplementary Materials

#### 1 Running time with respect to cluster size

| Cluster size | 10 | 20 | 30 | 40 | 50 | 60 |
|---|---|---|---|---|---|---|
| MUSCLE | 40.76 | 81.25 | 140.41 | 230.38 | 352.47 | 487.02 |
| Trellis BMA | 692.50 | 1360.38 | 2063.89 | 2678.58 | 3338.70 | 4027.76 |
| ITR | 175.36 | 729.61 | 1315.62 | 1150.22 | 1145.02 | 1151.44 |
| CPL | 6.81 | 25.67 | 60.98 | 69.76 | 70.14 | 69.77 |
| BBS | 1.51 | 1.84 | 2.13 | 2.48 | 2.71 | 3.07 |

Table 1: Average running time per cluster in milliseconds, with different cluster sizes. The experiments are done by running the algorithms on 1,000 clusters.

### 2 Performance of the tools on Simulated reads under IDS simulation

We tested the four algorithms on pure synthetic data, where the insertions, deletions, and substitutions happens uniformly at random at each position of the seed string, satisfying the IDS-channel assumption.

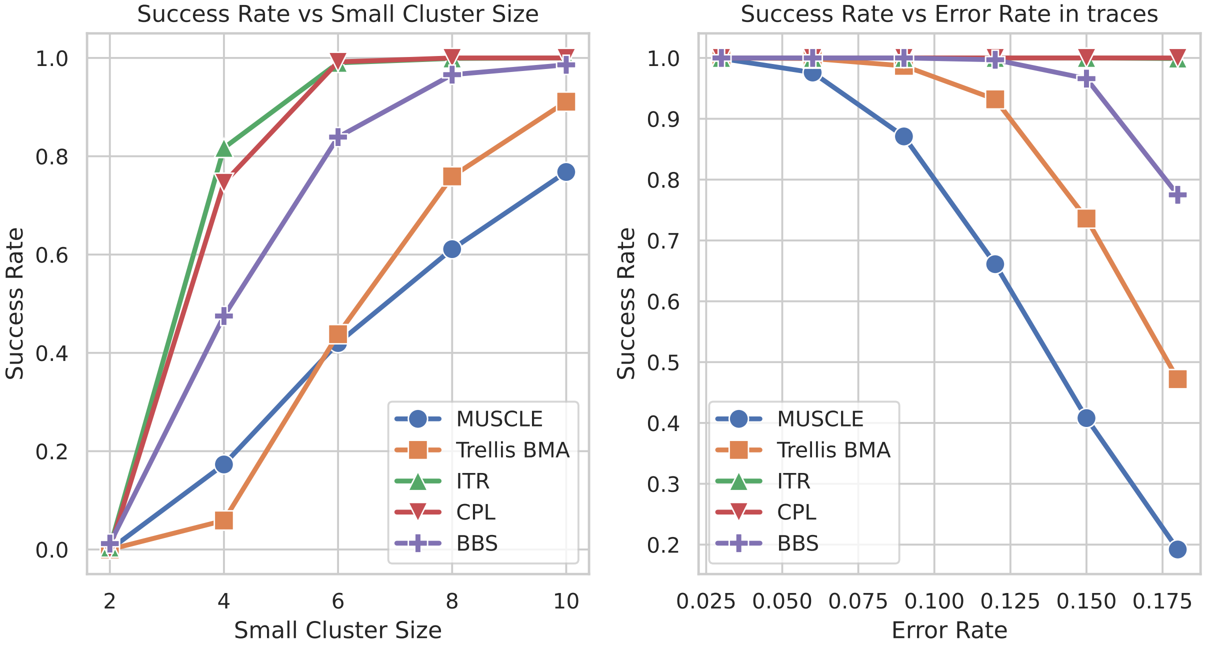


Figure 1: Performance of MUSCLE, Trellis BMA, ITR, and BBS on purely synthetic datasets satisfying the IDS-channel assumption. On the left plot, we fix the error rate of each trace to be 6%, and check the performance of the tools on very small cluster sizes. On the right, the cluster size is fixed to 20, and the tools are tested on synthetic datasets with different error rates, up to 18%.

We can notice that the ITR algorithm performs the best in both cases. Notably, it perfectly reconstructs all the clusters in the second experiment. The fact that BBS performs the best for simulated clusters using real data shows that the $k$-th order Markov Chain model is more flexible and accurate than the IDS-channel model in the real-life setting.

#### 3 Effect of beam width and $k$ on BBS

We tested the performance of BBS given different choices of parameters, the beam width $B$ and the order of the Markov Chain $k$, on the dataset of Srinivasavaradhan et al.

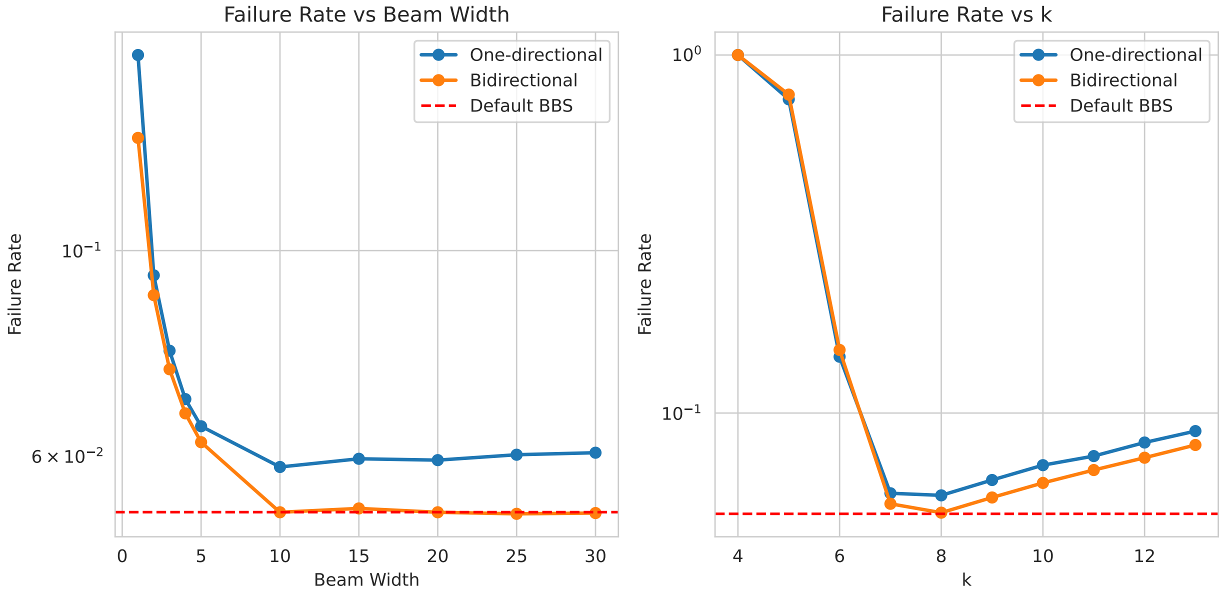


Figure 2: Performance of BBS on real datasets using various beam width (left) and fixed values of $k$ (right).

The performance of BBS showed a great increase for small $B$ from 1 to 5 and slowly stabilized for larger values than 20. There is even a slight decrease in performance for very large $B$, which might be because too many erroneous paths are included in the frontier during the search. In all cases, bidirectional beam search performed better than the usual one-directional beam search.

Next, we fix the beam width to be 20 and test the effect of $k$. Again, the performance increased significantly for small values of $k$ between 4 and 7, reaching the peak at $k = 8$, and gradually decreased for larger $k$. The low performance of $k$ is due to the uniqueness of $k$-mer being violated, creating loops in the De Bruijn graph, resulting in incorrect trace reconstruction. Larger $k$ than 8, on the other hand, encounter problems for high error rate and low coverage data as the correct $k$-mer doesn't always appear in the set of traces. The estimation of conditional probabilities is also less accurate due to the smaller sample size.

Notably, the default way of choosing $k$ by the BBS algorithm (that is, choosing the smallest $k$ such that all $k$-mers are unique in each trace) surpasses or matches the best-performing fixed values of $k$. The distribution of the chosen $k$ in the Microsoft dataset is as follows.

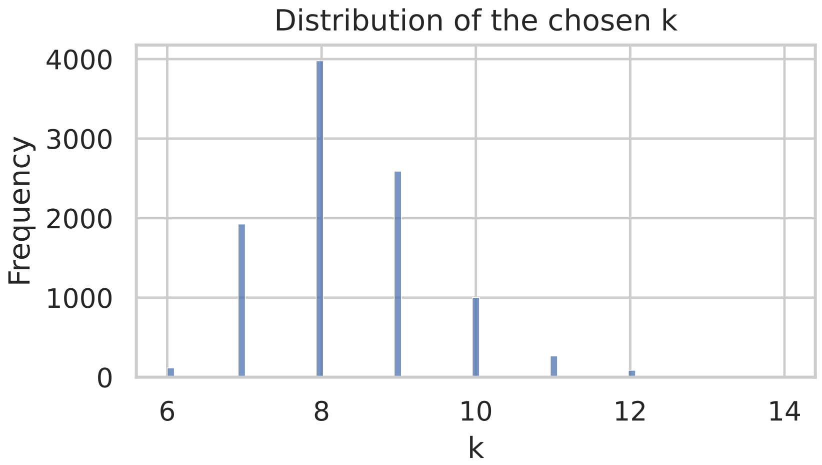


Figure 3: Distribution of the chosen $k$ values by the default BBS algorithm in the Microsoft dataset.

As expected, the time usage of the beam search algorithm doubles when performed in both directions (Figure 4). The time usage grows linearly with the beam width, with an exception at $B = 30$, which is because all the valid paths are already included in the frontier. We choose the default value of beam width to be $B = 20$, which appears to achieve great accuracy without sacrificing the speed.

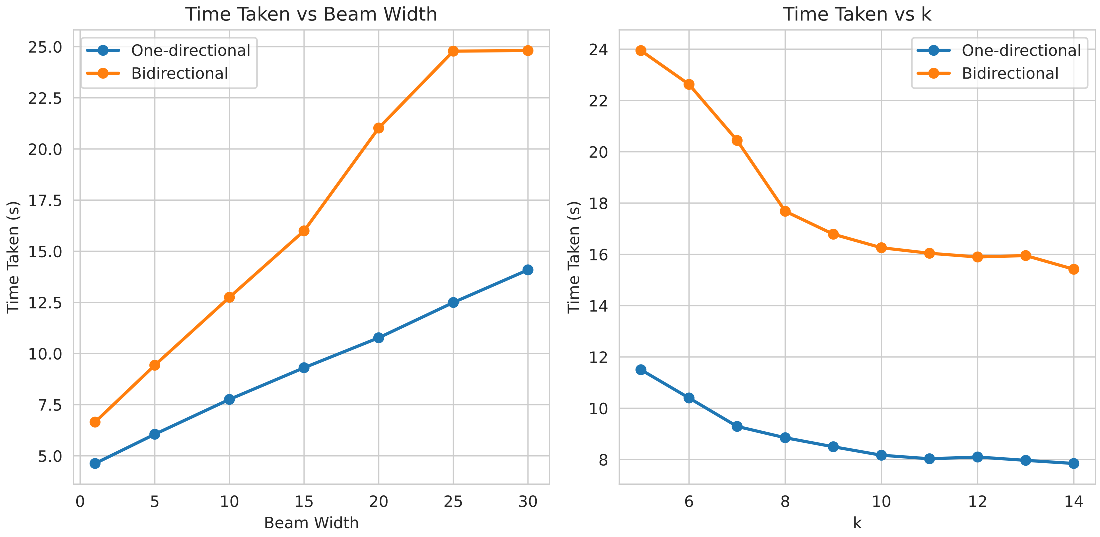


Figure 4: Time used to reconstruct all sequences in the Microsoft dataset (9964 clusters) given different beam width and fixed $k$.
